# β-glucan dependent shuttling of conidia from neutrophils to macrophages occurs during fungal infection establishment

**DOI:** 10.1101/512228

**Authors:** Vahid Pazhakh, Felix Ellett, Joanne A. O’Donnell, Luke Pase, Keith E. Schulze, R. Stefan Greulich, Constantino Carlos Reyes-Aldasoro, Ben A. Croker, Alex Andrianopoulos, Graham J. Lieschke

## Abstract

The initial host response to fungal pathogen invasion is critical to infection establishment and outcome. However, the diversity of leukocyte-pathogen interactions is only recently being appreciated. We describe a new form of inter-leukocyte conidial exchange called “shuttling”. In *Talaromyces marneffei* and *Aspergillus fumigatus* zebrafish *in vivo* infections, live imaging demonstrated conidia initially phagocytosed by neutrophils were transferred to macrophages. Shuttling is unidirectional, not a chance event, involves alterations of phagocyte mobility, inter-cellular tethering, and phagosome transfer. Shuttling kinetics were fungal species-specific, implicating a fungal determinant. β-glucan serves as a fungal-derived signal sufficient for shuttling. Murine phagocytes also shuttled *in vitro*. The impact of shuttling for microbiological outcomes of *in vivo* infections is difficult to specifically assess experimentally, but for these two pathogens, shuttling augments initial conidial redistribution away from fungicidal neutrophils into the favourable macrophage intracellular niche. Shuttling is a frequent host/pathogen interaction contributing to fungal infection establishment patterns.

## Introduction

In vertebrates, two phagocytic cell types have long been recognized as key players in the initial host defense response to infection: neutrophil granulocytes and macrophages [1]. Neutrophils and macrophages share many features: they are both migratory cells, they phagocytose microorganisms on encountering them, and they have intracellular mechanisms for killing microorganisms. However, although both phagocyte types engulf microorganisms, individual microorganisms interact with neutrophils and macrophages with different species-specific preferences, in different ways, and using different molecular mechanisms [2]. Conversely, the host has evolved diverse cellular strategies for these two different phagocytes to protect against the panoply of potentially pathogenic microorganisms.

The exchange of cytoplasmic material through contact-dependent mechanisms between adjacent cells is currently a topical field in cell biology. An example is the contact-dependent exchange of cytoplasm from macrophage to tumor cells as a metastasis-promoting mechanism [3], distinct from the cytoplasmic exchange between macrophages and tumor cells that occur via extracellular vesicles and nanotubes [4–6].

During infections, neutrophils and macrophages also engage in intercellular exchanges. Some microorganisms have evolved mechanisms that exploit these to enhance their pathogenicity and promote their spread between phagocytes. For example, *Yersinia pestis* and *Leishmania* promastigotes induce apoptosis in host neutrophils to then exploit efferocytosis, whereby clearance of dead neutrophils by macrophages leads to subsequent infection of this less hostile host cell [7–9]. Conversely, neutrophil phagocytosis of debris from dying macrophages is a recently-demonstrated method of mycobacterial dissemination [10]. *Candida albicans* [11] and *Cryptococcus neoformans* [12] can be ejected from host macrophages by non-lytic exocytosis, while macrophage-resident *C. neoformans* [13] and *Aspergillus fumigatus* [14] also can enter new host macrophages through lateral transfer (recently termed metaforosis [15]). The Gram negative bacteria *Francisella tularensi* and *Salmonella enterica* are transferred between macrophages by a process related to trogocytosis [16]. These scenarios are characterised either by death of the donor cell, expulsion of the pathogen from the donor cell without direct contact between donor and recipient phagocyte, or transfer between the same type of phagocyte. None involves transfer by direct contact from a living neutrophil to living macrophage.

Such interactions provide an opportunity for intracellular pathogens to transfer to a new host cell, while minimising exposure to a potentially hostile extracellular environment. As antibiotic resistance becomes a growing problem, there is an ever-increasing interest in host-pathway directed anti-infective therapies. Host dependent processes for pathogen dissemination represent key potential targets [17].

Zebrafish have emerged as an ideal model for intravital imaging of leukocyte behaviors during infection [18]. They combine the advantages of small size, optical transparency (particularly as embryos and larvae) and suitability for genetic manipulation. Zebrafish phagocytes have been comprehensively characterized in developmental, genetic and functional studies [19].

Our recent modeling of fungal infections in zebrafish models have focused on high spatiotemporal resolution intravital imaging of the initial leukocyte-pathogen interactions [20]. During these studies, we observed a form of microorganism exchange between neutrophils and macrophages that we believe to be previously undescribed, which we have named “shuttling”. In shuttling, a living donor neutrophil laden with previously-phagocytosed fungal spore(s) transfers this cargo to a recipient macrophage through a tethered direct contact, without death of the donor neutrophil. Shuttling is therefore different to all the previously described microorganism exchanges between phagocytes.

In the present study, we comprehensively describe neutrophil-to-macrophage “shuttling”. Studying shuttles presented considerable technical challenges, as they could only be identified by directly observing them retrospectively in *in vivo* live imaging datasets. To recognize a shuttle, all three phases of the process had to be captured in the imaged volume: initial carriage of a phagocytosed spore within a mobile, living donor neutrophil; the moment of intercellular contact and transfer between neutrophil and macrophage; and the departure of the previously unladen recipient macrophage, now carrying its newly acquired cargo. All three shuttle-defining steps needed to have occurred within the imaged volume, despite the high mobility of the participating cells. Despite this challenge, we comprehensively describe the morphology of shuttling, quantify key parameters of the dynamic transfer process, and identify a key mechanistic determinant by demonstrating that the conidial cell wall component β-glucan is a fungal-derived molecular signal sufficient to trigger shuttling of particles. Additionally, by replicating this phenomenon using murine phagocytes *in vitro*, we provide evidence that shuttling is a conserved behavior of both fish and mammalian phagocytes.

## Results

### Some *Talaromyces marneffei* conidia phagocytosed by neutrophils are “shuttled” to macrophages

While studying leukocyte behavior during the establishment of *Talaromyces marneffei* infection following inoculation of live conidia into zebrafish [20], we unexpectedly observed the recurrent direct transfer of conidia from live neutrophils to adjacent live macrophages (Fig 1, Supplementary Movie S1a,b). The phenomenon was revealed by combining a 3-color fluorescent reporter system (labelling neutrophils in green [EGFP], macrophages in red [mCherry], conidia in blue [calcofluor]) with high spatiotemporal resolution live confocal imaging. We called this new form of inter-phagocyte pathogen transfer “shuttling”. Two defining features of shuttling distinguished it from other previously-described forms of pathogen transfer. Firstly, shuttling occurred between live leukocytes, demonstrated by the mobility of both donor neutrophil and recipient macrophage before, during and after shuttles. Secondly, the dynamic morphology of shuttling suggested purposeful rather than random exchange, through a tethered cell-to-cell contact.

**Figure 1.**
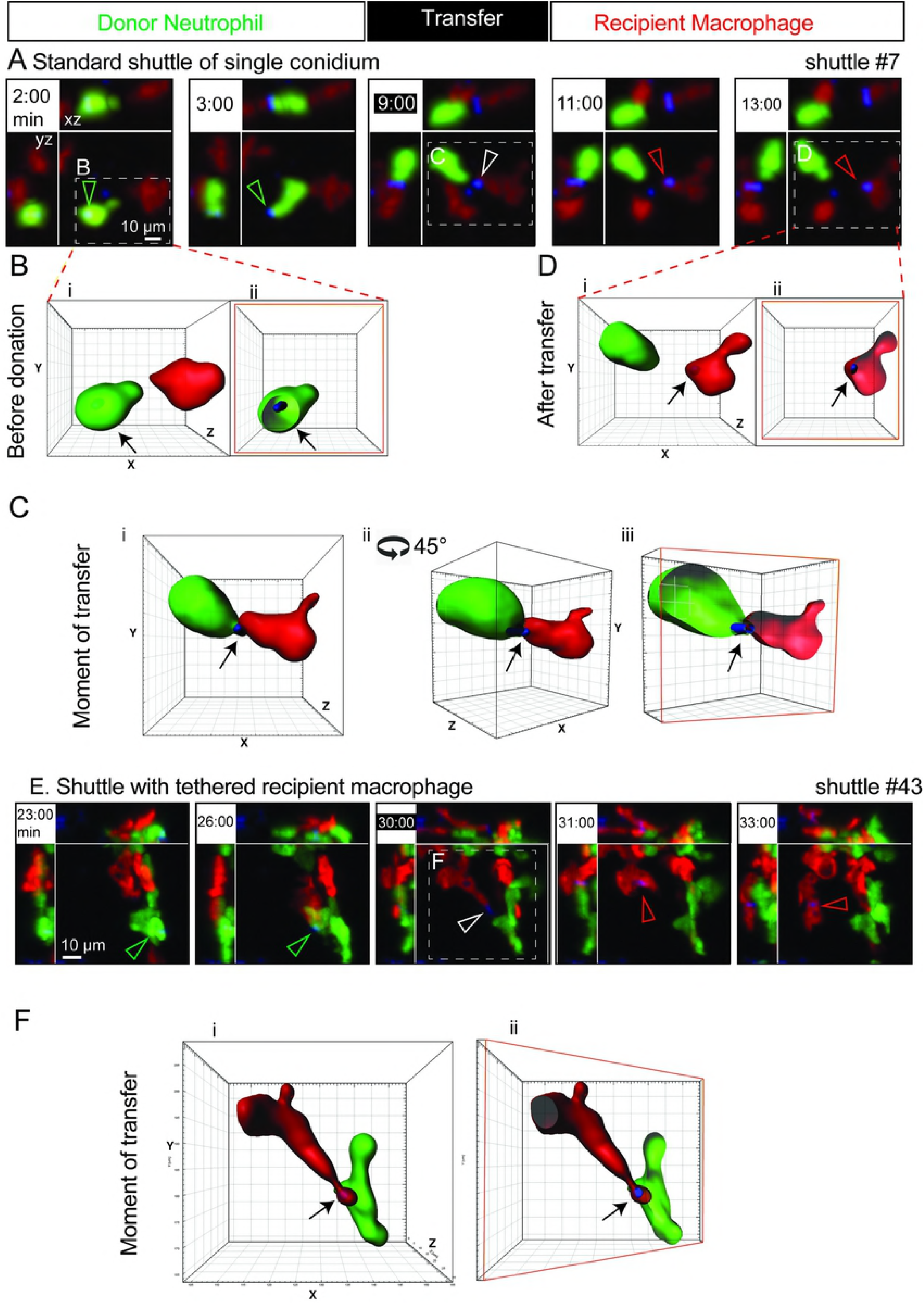
Shuttling of individual *T. marneffei* conidia from neutrophil to macrophage. A. A representative standard shuttle of a calcofluor-stained conidium (blue) from a *Tg(mpx:EGFP)* neutrophil (green) to a *Tg(mpeg1:Gal4FF)x(UAS-E1b:Eco.NfsB-mCherry)* macrophage (red), corresponding to the example in Supplementary Movie S1a. Panels include isometric orthogonal yz and xz views corresponding to the xy maximal intensity projection, and indicate the time in minutes from start of movie. The time point colored white-on-black is the moment of transfer. Colored arrowheads indicate the conidium within donor neutrophil (green), at the point of intercellular transfer (white) and in the recipient macrophage (red). B-D. Volume-rendered views of the standard shuttle in (A), detailed before (B), at the moment of transfer (C) and after (D), demonstrating the intracellular location of the shuttled spore in donor neutrophil and recipient macrophage, the focal intercellular contact at the moment of transfer. Cii is the image in Ci rotated 45° around a central vertical axis in the direction shown. Images Bii, Ciii and Dii are sectioned volume-rendered views; a sectioned plane is represented by a red box. E-F. Shuttle demonstrating tethering of the departing recipient macrophage following a shuttle. Panel E presentation organized as in panel A. The tethered moment of transfer is detailed by volume-rendering in F, presented as in panels B-D. Scales as shown. Stills in A and E correspond to Supplementary Movie 1a,b respectively.

To characterize the dynamic morphology of shuttling comprehensively, we systematically collected multiple unselected examples from extensive confocal live-imaging microscopy experiments. To ensure that shuttles were unequivocally distinguished from all other modes of intercellular pathogen transfer, stringent criteria were applied for events to be included in this initial panel. For inclusion as a shuttle, all three phases of donation, transfer and receipt were required to be unequivocally visualized (see Materials and Methods for full details). The resulting collection of unequivocal shuttles comprised 13 examples of live *T. marneffei* conidial shuttling (Fig S1, Supplementary Figure S1), and as shuttling mechanisms were explored, another 17 unequivocal examples of conidial shuttling and 18 examples of the shuttling of other particles meeting all stringent definition criteria were collected (Supplementary Table S1).

Shuttling of live *T. marneffei* conidia occurred only in the first two hours of infection establishment (median time of shuttle, 33 min [range 14-97] from commencement of imaging; n=13 shuttles collected in 69 movies; Supplementary Fig S1A). In contrast, no *T. marneffei* shuttles occurred during >181 hr of imaging after 2 hr post inoculation.

These *T. marneffei* shuttling examples exhibited morphological features with mechanistic implications. In several cases, the donor neutrophil and/or recipient macrophage formed a highly polarised shape resulting from cell-to-cell tethering around the time of shuttling (Fig 1E,F, Fig 2A, Supplementary Movie S1b-d). These drawn-out tethered extensions of neutrophil and macrophage cytoplasm before, during or after shuttling indicated a focal rather than a whole-of cell “hugging” interaction between them. Furthermore, this tethering confirms that the cells come into direct physical contact for the shuttle, rather than merely moving into close proximity and transferring the conidium by expulsion into the extracellular space and re-phagocytosis.

**Figure 2.**
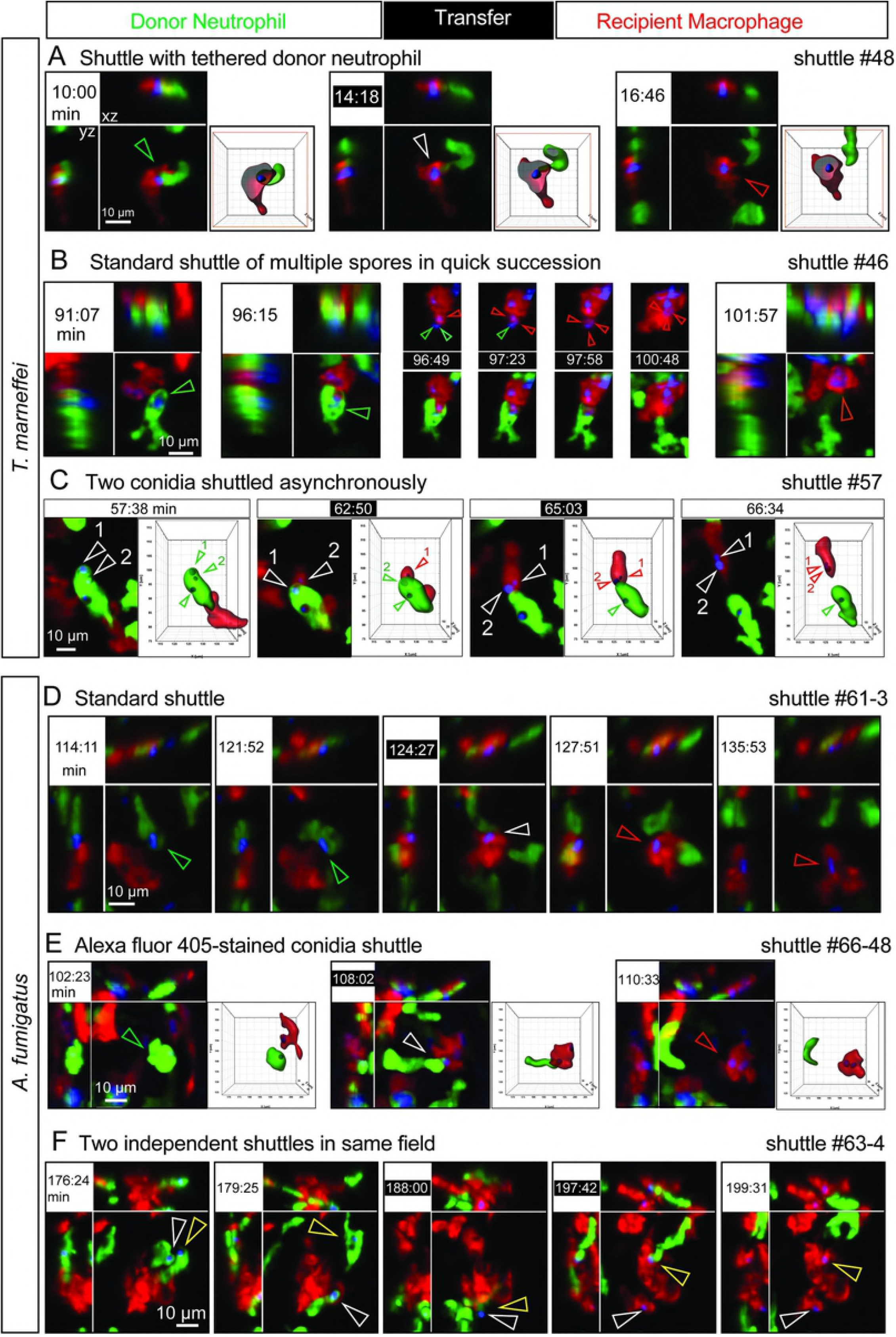
Variant shuttles of fungal conidia from neutrophil to macrophage. A variety of shuttles of conidia (blue) from *Tg(mpx:EGFP)* neutrophils (green) to *Tg(mpeg1:Gal4FF)x(UAS-E1b:Eco.NfsB-mCherry)* macrophages (red). In each example, panels include isometric orthogonal yz and xz views corresponding to the xy maximal intensity projection, and indicate the time in minutes from start of movie. Time points colored white-on-black are the moments of transfer). (A,C,E) include volume-rendered views corresponding to the maximal intensity projection; where this volume is sectioned, the framing box is shown in red. Colored arrowheads indicate the conidium within donor neutrophil (green), at the point of intercellular transfer (white) and in the recipient macrophage (red). **A-C. Shuttles of *T. marneffei* conidia.** A. Shuttle demonstrating tethering of the donor neutrophil at the moment of transfer. B. Shuttle of multiple conidia from one donor neutrophil in quick succession. First two frames show donor neutrophil laden with multiple conidia at two pre-shuttle time points. Frames from t=96:49-100:48 sec are maximal intensity projections only, encompassing the shuttling transfer of 3 conidia over 4 minutes. Upper row of panels shows red (macrophage) and blue (conidia) channels only, lower row of panels includes the green channel (donor neutrophil). C. Shuttle of two conidia from one donor neutrophil one after the other at an interval of 2 min 13 seconds. Volume-rendered images corresponding to maximal intensity projections show the two shuttled conidia (labelled 1 and 2) before, during and after the shuttle. **D-F Shuttles of *A. fumigatus* conidia.** D. Standard shuttle of single conidium. E. Standard shuttle of conidium labelled with Alexfluor 405 rather than calcofluor, accompanied by volume-rendered images which focus attention onto the conidium and donor neutrophil of interest. F. Two independent shuttles by different donor neutrophils occurring in the same field. In this series, the course of each shuttled spore is followed by white and yellow arrowheads. Scales as shown. Stills in A-E correspond to Supplementary Movies S1c, e, f, and S2a, c, e respectively.

Although single conidia were usually shuttled (Fig 1, Supplementary Table S1), occasionally more than one conidium was transferred (2/13 instances; Fig 2B,C, Supplementary Movie S1e-f). One example of this was in quick succession (Fig 2B, Supplementary Movie S1e). However, the non-synchronous transfer of two shuttled conidia in series from the same donor neutrophil in another example (Fig 2C, Supplementary Movie S1f) indicated that the signalling mechanism driving each shuttle could operate independently.

### Shuttling also occurs with *Aspergillus fumigatus* conidia

To test whether conidial shuttling was specific to *T. marneffei* or a general phenomenon of fungal infection establishment, we assayed for shuttling following inoculation with live *Aspergillus fumigatus* conidia, another fungus whose interactions with leukocytes are well studied in zebrafish models [20–23], but for which shuttling has not previously been described. Seven unequivocal shuttles of live *A. fumigatus* conidia occurred in 6/22 imaging sequences (Fig 2D-F, Supplementary Fig S1B, Supplementary Movie S2). The median time of shuttling was 121 [range 30-199] min following commencement of imaging. *A. fumigatus* shuttles exhibited similar features to *T. marneffei* shuttles, including cell-to-cell tethering (Supplementary Movie S2d). In 1/7 example, two shuttles occurred in the same imaged volume between different donor neutrophils and macrophages, separated by an interval of 10 minutes (Fig 2F).

To exclude the possibility that shuttling was an artefact of labelling conidia with calcofluor, we tested conidia with an alternate label. *A. fumigatus* conidia labelled with Alexa Fluor^®^ 405 were also shuttled (Fig 2E, Supplementary Movie 2c). All shuttles occurred from donor neutrophil to recipient macrophage. No macrophage-to-neutrophil shuttles were observed despite looking carefully for them.

Collectively, these observations establish that shuttling is a recurrent form of unidirectional pathogen transfer from neutrophils to macrophages that occurs early in fungal infection establishment. It is not a peculiarity of the host response to a particular fungal pathogen, because it occurs with two fungal species.

### Incidence of shuttling

Shuttling events meeting our stringent criteria were observed in 20/91 (22%) unselected imaging sequences of >60 min duration (Supplementary Fig S1). While this ascertainment rate provided a scorable surrogate categorical variable for shuttling incidence, a more biologically-relevant measure of the incidence of shuttling would add weight to its biological significance.

One such biologically-relevant quantification is the shuttling incidence per condium *at risk of shuttling*. This measure, regardless of any macrophage phagocytic activity and recruitment of non-phagocytosing leukocytes, is denominated solely by the number of spore-laden neutrophils in the imaged volume available to act as donors. While this is impossible to determine exactly for any single image series due to neutrophil flux through the imaged volume, it is possible to compute an averaged estimate. For both these fungi, we have previously examined phagocytosis during infection establishment and previously reported that macrophage phagocytosis predominates over neutrophil phagocytosis in the first three hours following inoculation [20]. These phagocytosis data resulted from analysis of a subset of imaging files of the current dataset, and so provide a basis for estimating an averaged shuttling incidence based on averaged spore-laden neutrophil phagocytosis rates. For *T. marneffei*, an average of 1.34 conidial-loaded neutrophils (67 neutrophils at 50 time points) were present at any time in the imaged volume to be available as donors throughout the first 180 minutes after inoculation (derived from n=10 imaging series, being those 10 series closest to 180 min in length). Hence 13 shuttles in 69 imaging series means that on average, 14% of spores available in neutrophils for donation were shuttled in 3 hours. Also of note is the fact that 5/10 imaging series had only ≤1 spore-laden neutrophil present in the imaged volume during the first 180 minutes, hence these imaging series provided little opportunity for shuttling to occur. For *A. fumigatus*, neutrophil phagocytosis of conidia was much rarer, as also observed by others [20,24]. An analogous averaged calculation gives an average incidence of 44% for shuttling of *A. fumigatus* spores that were available for donation by loaded neutrophils in the 3 hours after inoculation (36 neutrophils / 50 time points = 0.72 spore-laden neutrophils on average at any time point over three hours; 7 shuttles in 22 movies).

Calculating the real shuttle incidence is challenging and these rates are certainly underestimates. The rate depends on the sensitivity of ascertainment, which is constrained by the limitations of the detection method and our stringent definition of shuttling. These two factors together conspire to underestimate shuttle incidence.

Challenges in shuttle detection that contributed to underestimating incidence included: (1) shuttles can currently only be recognised by the laborious method of manually observing them in retrospective analysis of imaging datasets; (2) at the magnification required for the sub-cellular resolution needed to see shuttles, the imaged volume is only a small fraction of the infected volume; and (3) the leukocytes involved are highly mobile and frequently move out of the imaged volume, hence the denominators for computing incidence are constantly changing.

Several other observations indicate that shuttles are not rare. If shuttles were rare, it would be unlikely that multiple examples would occur together or in the same imaging sequence. However, 5/20 datasets contained examples of multiple shuttles, either 2-3 spores being shuttled together, or in quick succession, or asynchronously from the same or several different donor neutrophils (Fig 2B,C,F, Supplementary Movies S1e,f, S2e; Fig 3, Supplementary Movies 2f, 3a-c).

**Figure 3.**
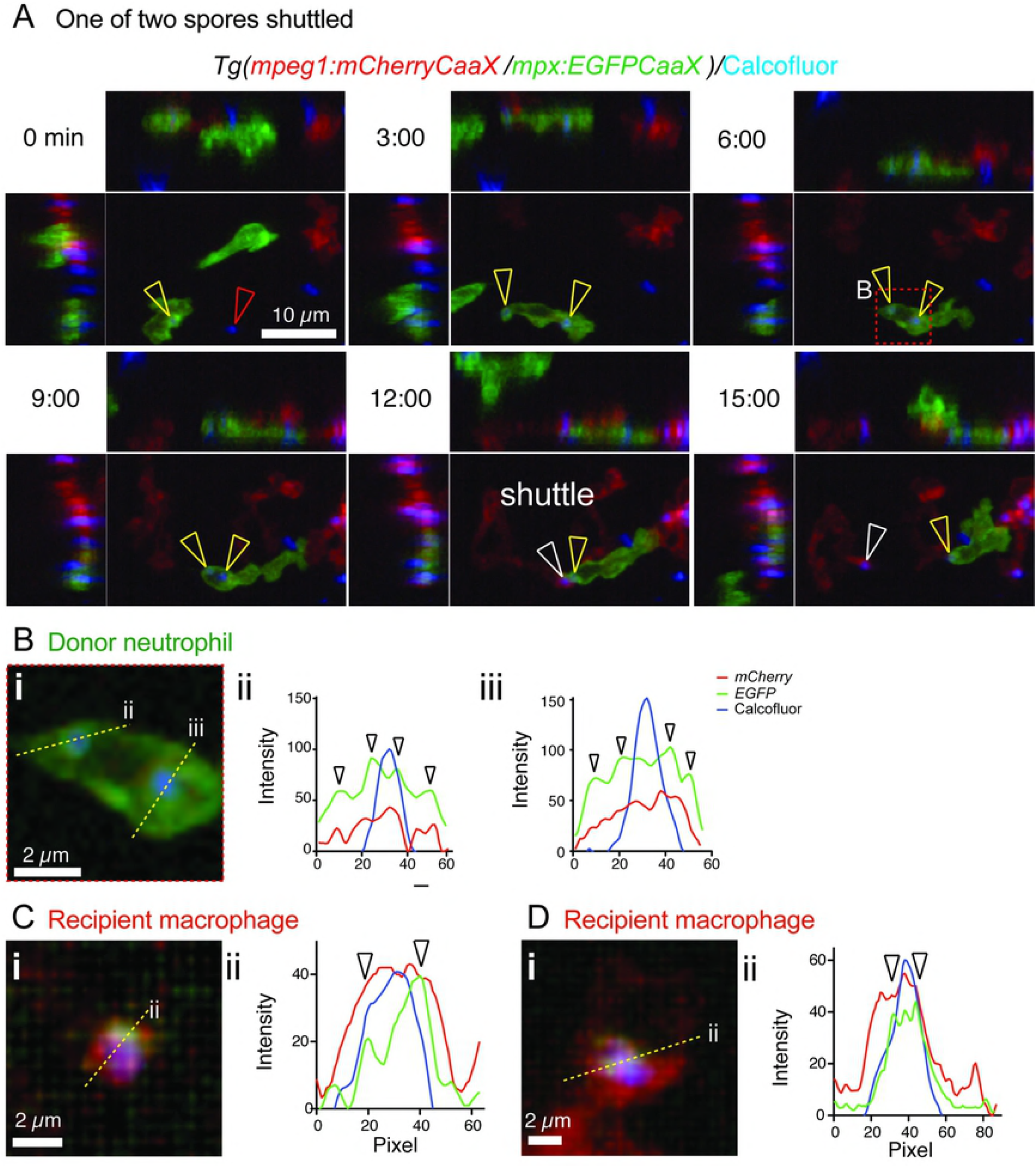
Shuttling of *T. marneffei* conidia between neutrophils and macrophages involves phagosome transfer. A. Shuttle of calcofluor-stained conidium (blue) from *Tg(mpx:EGFPCaaX)* neutrophils (green) to *Tg(mpeg1:mCherryCaaX)* macrophages (red). These reporter lines have membrane-localised fluorophore expression. Panels include isometric orthogonal yz and xz views corresponding to the xy maximal intensity projection, and indicate the time in minutes from start of the movie. Colored arrowheads indicate a conidium before it is phagocytosed by the donor neutrophil (red), conidia within the donor B. (i) Detail of the boxed area of the donor neutrophil in (A, 6 min panel). Yellow dotted line indicates the position of the cross-section for the 3-color fluorescence intensity plots in (ii) and (iii). Both shuttled and non-shuttled conidia are flanked by peaks of green fluorescence, consistent with their location in a membrane-lined phagosome. C,D. Cross-sections fluorescence intensity profiles (ii) corresponding to the yellow lines in (i), for two macrophages that received a spore from a neutrophil in this dataset, which contained 3 independent spore shuttles. The arrowed EGFP-channel signal demonstrates the transfer of neutrophil-derived EGFP-tagged membrane in the vicinity of the spore (blue channel signal). Scales as shown. Stills in A correspond to Supplementary Movie 3a.

The stringent criteria applied to ensure only unequivocal shuttles were included also means that the shuttle incidence is likely to be underestimated. Multiple events that were probably shuttles were excluded from this initial panel of unequivocal shuttles (see examples in Supplementary Figure S2, Supplementary Movie S4). There were probable shuttles where the conidium could not unequivocally be resolved as within the donor neutrophil rather than adherent to it (Supplementary Figure S2A, Supplementary Movie S4a). The criterion most often not met was clear visualisation of the donor-recipient cell-to-cell contact at the point of conidial transfer (Supplementary Figure S2C-D, Supplementary Movie S4c-d). This scenario included instances where a phagocytosed particle appeared to be deposited by the neutrophil into extracellular space and was then subsequently taken up by a macrophage. In some imaged volumes, a large number of highly active neutrophils and macrophages were attracted to the inoculated spores, and it was impossible to separate what happening although initially many spores were in neutrophils that ended up in macrophages (Supplementary Figure S2D, Supplementary Movie S4d). These imaging series have been included in the denominator of our unselected series.

From these data and considerations, we conclude that although the detection of shuttling is laborious and challenging, shuttling itself is not a rare phenomenon. For both these fungal pathogens, those spores that are initially phagocytosed by neutrophils have a substantial chance of being shuttled to macrophages in the first 3 hours of infection establishment.

### Shuttling involves phagosome transfer

We previously reported the transfer of neutrophil cytoplasm to macrophages in the context of inflammation [25]. We therefore hypothesised that shuttling could also involve transfer of donor neutrophil cytoplasmic components to the recipient macrophage. To test specifically whether neutrophil membrane was also transferred, we imaged shuttling in *Tg(mpeg1:mCherry-CaaX/mpx:EGFP-CaaX)* embryos, in which the fluorescent labelling of neutrophils and macrophages is localised to the membrane via a prenylation motif (Fig 3, Supplementary Movie S3). Cross-sectional fluorescence intensity profiling of conidia in these transgenic lines demonstrated that prior to shuttling, about-to-be shuttled conidia reside within membrane-bound compartments within the neutrophil (Fig 3B). Observation of shuttled conidia in macrophages immediately following transfer demonstrated that green fluorescent signal surrounding the spore attributable to neutrophil membrane was also transferred to the macrophage (Fig 3C).

This provides direct evidence that shuttled conidia are located in a membrane-lined sub-cellular neutrophil compartment, likely to be a neutrophil phagosome, which is shuttled in its entirety to the recipient macrophage. The rapid decay of the cytoplasmic neutrophil reporter fluorophore signal following shuttling suggests that within the macrophage it is either quenched due to pH change, or that the structure of the shuttled phagosome and its component proteins are rapidly destroyed by the macrophage.

### Phagocyte motility confirms that living cells participate in shuttling

Our previous studies demonstrated that phagocytes exhibit lineage and site-specific spatiotemporal responses during establishment of fungal infection [20]. We asked whether shuttling occurred “on the fly” between fast moving cells, or if cells slowed down and “parked” to engage in this intercellular interaction.

We first focussed on the scenario in which *T. marneffei* conidia were delivered into the somite. To characterize the overall picture of leukocyte movement in which shuttling occurred, we used four-dimensional cell tracking in Imaris software (Bitplane) to extract and plot cell coordinates in time and space. We interrogated these data using the open source programming language R (Fig 4A), as has been used to analyse leukocyte swarming [26]. In this scenario, neutrophils started to migrate towards the infection site soon after inoculation with conidia, while macrophage migration initiated later, during the second hour post infection. Overall, neutrophils exhibited more rapid motility than macrophages at all times and in all directions (P<0.0001). Phagocytosis of conidia upon arrival at the site of infection was associated with a reduction in migration velocity for both neutrophils and macrophages (Fig 4A).

**Figure 4.**
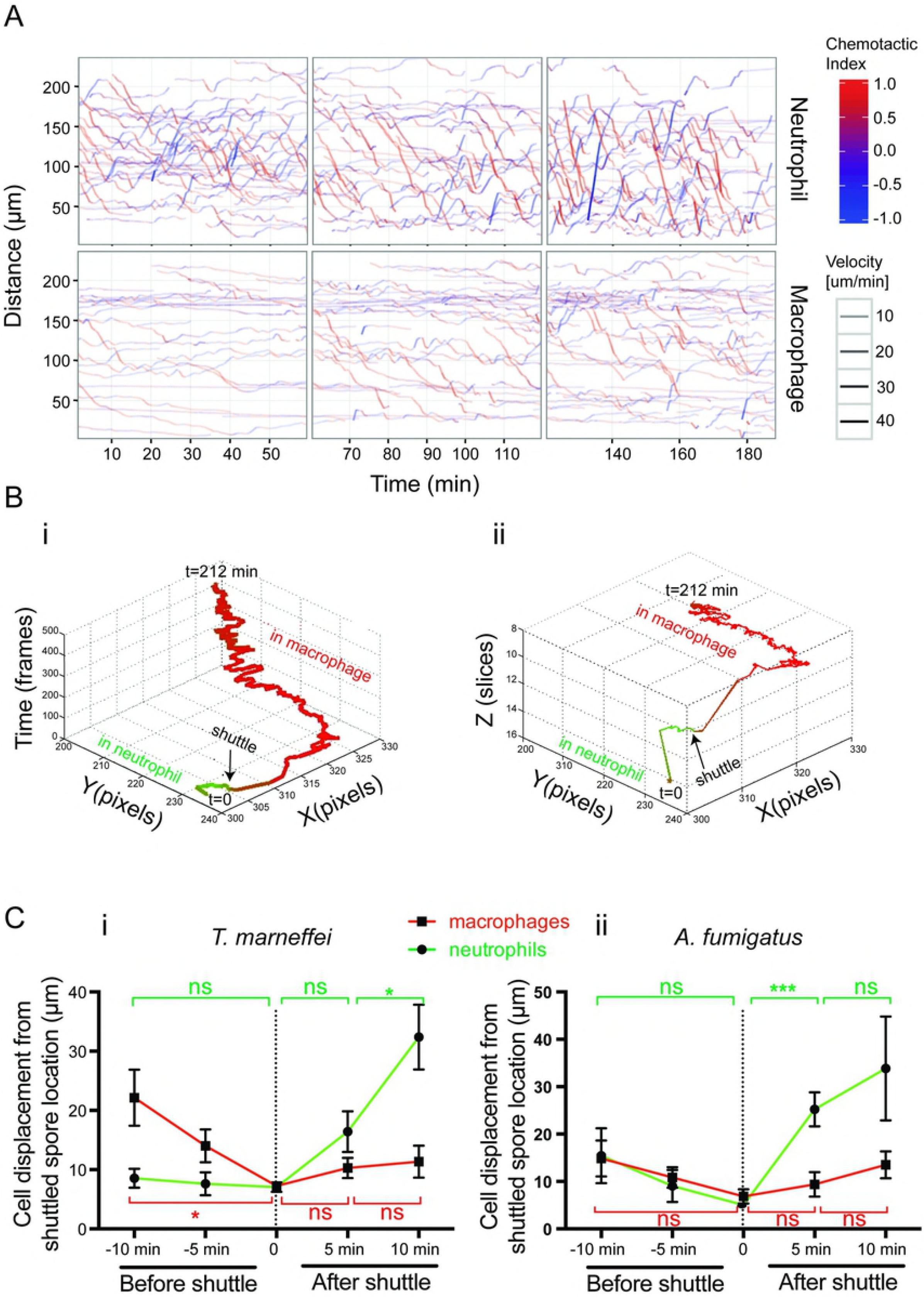
Phagocyte mobility during shuttling. A. Cell tracking analysis of neutrophil and macrophages following intramuscular inoculation of *T. marneffei* conidia. A chemotactic index (red, movement towards infection; blue, movement away) and velocity (thickness of the line). Neutrophil migration towards the infection is earlier and faster; macrophage migration is later and slower. B. Output of ShuttleFinder software, for the shuttle shown in Fig 1A. Track color indicates the cellular context of the conidium (green, neutrophil; red, macrophage). (i) shows the shuttled spore track in xy dimensions and time. (ii) shows it in xyz dimensions. The two outputs collectively show that the shuttle occurred between donor and recipient phagocytes that were mobile in both space and time. Imaging parameters: x, y, 1.31 pixels/μm; z interval, 3.3 μm/slice; frame rate, 28.0 sec/frame. C. Plots of donor neutrophil and recipient macrophage cell displacement from the shuttle spore location over two fixed 5 min time intervals before and after the moment of shuttle, for *T. marneffei* shuttles (i), and *A. fumigatus* (ii). Note that displacement is a measured distance without directional information. Data are mean±SEM at each time point (i, n=10; ii, n=7). P-values from paired t-tests: * <0.05; ***≤0.001; ns, not significant.

To focus on the subset of phagocytes engaging in shuttling within this melee of phagocyte activity, we developed “ShuttleFinder”, a Matlab^®^ program based on “PhagoSight” [27] that performs spatiotemporal tracking of conidia and reports the colour of their immediate surrounding environment (Fig 4B). ShuttleFinder did not facilitate automatic shuttle discovery due to the high number of disjointed tracks and a high number of false positives. However, it enabled the paths of conidia that were shuttled to be displayed in 2-dimensional space and time (Fig 4Bi) and 3-dimensional space (Fig 4Bii). This demonstration of conidial translocation showed the extent to which the donor neutrophils and recipient macrophages in which shuttled spores resided were mobile prior to and following the shuttle (Fig 4B,C).

We next assessed if cell velocity changed during shuttling manually examining the displacement of neutrophils and macrophages over fixed periods (5 and 10 minutes) before and after shuttles. This analysis revealed that in the 10 minutes prior to shuttling, the displacement of a donor neutrophil from the shuttle location was <15 μm (i.e. < 2 cell diameters), whereas after shuttling, neutrophil displacement significantly increased, indicating neutrophil velocity was significantly faster after shuttling than before it. This occurred for both *T. marneffei* and *A. fumigatus* shuttles (Fig 4C). The higher velocity of neutrophils after shuttling strongly indicates that the donor neutrophil was alive.

Macrophages also moved toward the shuttle point, their displacement altering significantly only in the case of *T. marneffei* shuttles. Macrophage displacement did not alter significantly in the 10 minutes after receiving a shuttled spore (Fig 4C). However, over longer periods of time, the recipient macrophages were also mobile (Fig 4B), confirming their viability and indicating that receiving a donated spore is accompanied by only a temporary reduction in migratory activity.

The 20 shuttles of live spores meeting all stringent definition criteria (Fig 5A) suggested that shuttling continues the process of conidial dissemination. In 16/20 cases the donor neutrophil entered the field during the imaging period, and hence it had picked up its spore for donation elsewhere. Furthermore, in 18/20 cases, the recipient macrophage separated from the donor neutrophil before the image sequence finished (Fig 5A).

**Figure 5.**
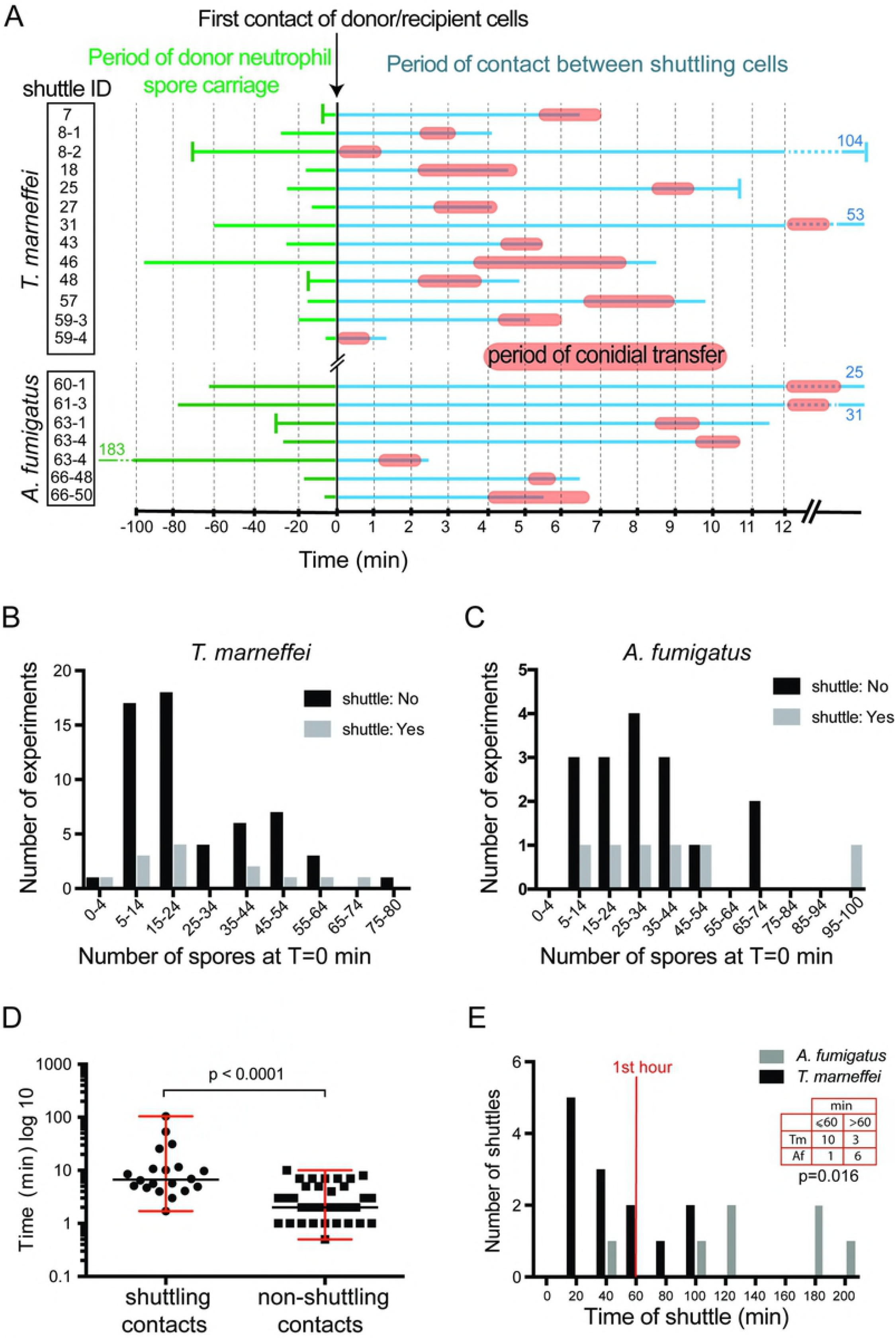
Dynamics of neutrophil to macrophage conidial shuttles. A. Time maps of cellular contact during shuttles. Charts are aligned with time=0 at the point of initial donor/recipient cell contact. Chart shows: time of conidial residence in donor neutrophil prior to intercellular contact (green line; vertical bar indicates resident at start of imaging; otherwise line start indicates point of neutrophil phagocytosis); time of general intercellular contact (blue line); period of contact during actual conidial transfer (orange bars). Charts are for each of 20 stringently-defined unequivocal shuttles (n=13 *T. marneffei*, n=7 *A. fumigatus*) tabulated and identified as in Supplementary Figure S1. The entire sequence from phagocytosis to cell separation is shown unless vertical bars at beginning or end of lines indicate the start or end of the imaging file; blue and green numbers indicate endpoint times where lines have been clipped. B, C. For *T. marneffei* (B) and *A. fumigatus* (C) infections, histograms of number of experiments with and without observed shuttles, by number of conidia present in the initial imaging volume. D. For conidia-laden neutrophils, duration of shuttling contacts (n=20 shuttles) compared to random non-shuttling contact (n=34). Data are summarized by median and range. E. Distribution of times of shuttle after imaging commenced for *T. marneffei* and *A. fumigatus*. P=0.016 for categorical variable of shuttles occurring at ≤60 and >60 min between the two fungal species (Fisher’s exact test).

In summary, these data show that living neutrophils slow down to donate conidia for shuttling, that the living recipient macrophages are relatively stationary at the time of transfer and temporarily parked following receipt of a shuttled spore, and that shuttling contributes to the ongoing process of spore dissemination during infection establishment.

### Shuttling is a purposeful interaction that involves sustained intercellular communication

We considered the possibility shuttling might be a chance event rather than purposeful interaction. If shuttling occurred by chance alone, then the number of shuttles would be expected to be a function of the number of conidia delivered. However, shuttles occurred in fields that had initially as few as <4 conidia, and as many as 75-100 conidia, for both fungal species, and we could not resolve a trend based on the number of conidia initially in the imaged volume (Fig 5B,C).

We also hypothesized that if shuttling were purposeful rather than a random event, then this would be reflected by purposeful intercellular interactions. Shuttling is characterized by multiple polarized interactions between the donor neutrophil and the recipient macrophage prior to the shuttle, indicating sustained and purposeful prior cell-cell communication (Supplementary Movies S1-3). To quantitatively test the hypothesis that the degree of intercellular interaction between shuttling leukocytes was unusually extensive, we compared the duration neutrophil/macrophage contacts that ended with a shuttle to randomly selected non-shuttling contacts. This analysis revealed that shuttling cells stay in contact for a significantly longer period prior to shuttling than is otherwise the case for random neutrophil/macrophage interactions (P<0.0001) (Fig 5D). This observation is consistent with there being signals bringing the donor and recipient cells together prior to shuttling. Furthermore, it indicates that these signals are different from those that attracted the leukocytes to migrate to the site of infection.

### Pathogen-dependent shuttling kinetics indicate a conidial determinant

We hypothesised that the molecular mechanism driving shuttling likely involved fungal determinants. If this were the case, this might result in different kinetics for the shuttling of conidia of different fungal species. Although we observed no obvious morphological difference between shuttles of *T. marneffei* and *A. fumigatus* conidia (suggesting that the mechanism driving shuttling is fundamentally the same for both), the kinetics of shuttling events differed for the two species. *T. marneffei* shuttles occurred predominantly in the first hour after inoculation, whereas *A. fumigatus* shuttles happened at later time points (p=0.016) (Fig 5E). There was no ascertainment bias for shuttles of one or other fungus, as the two movie datasets shared a similar distribution of imaging durations (Supplementary Fig S1).

This important observation indicates that although shuttling is a general phenomenon in fungal infection establishment, because the kinetics of shuttling is specific to the fungal species, determinants of the molecular mechanism reside in properties of the conidia themselves.

### The fungal determinant of shuttling is not a metabolic product

A fungal determinant of shuttling could be either a chemical constituent of the conidium, or a newly-synthesized metabolite of the germinating fungi. To distinguish between these possibilities, we microinjected conidia inactivated by either freezing (*T. marneffei*) or γ-irradiation (*A. fumigatus*) and imaged for shuttling events. We observed multiple shuttles of inactivated fungal conidia (Figs 3, 6A; Supplementary Movie S3), confirming that shuttling was stimulated by non-temperature-labile components of the conidial cell wall, rather than an actively synthesized signal.

**Figure 6.**
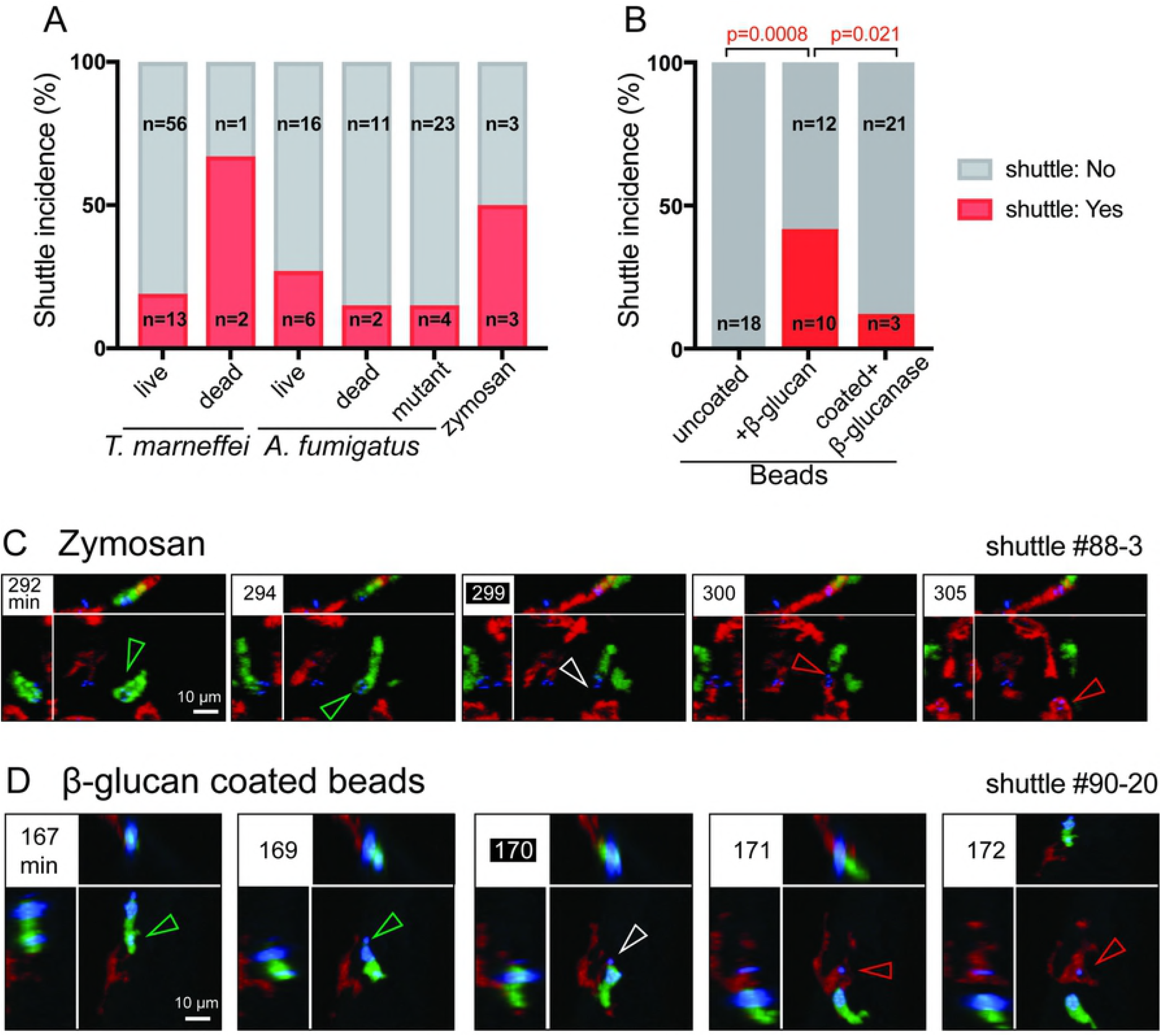
β-glucan is a fungal determinant sufficient to trigger shuttling. A,B. Relative frequency of shuttles for different cargoes, incidence computed for each condition as number of 3 hour imaging datasets with shuttle(s) / total number of imaging datasets. By Chi-squared analysis, there are no significant differences for the comparisons: live spores of the two species (p=0.38); live and dead *T. marneffei* (p=0.31); dead spores of the two species (p=0.34). n values indicate the number of datasets in each category. C,D. Images of representative shuttles of zymosan particle (C) and β-glucan coated plastic beads (D). Shuttles of particles (blue) are from *Tg(mpx:EGFP)* neutrophils (green) to *Tg(mpeg1:Gal4FF)x(UAS-E1b:Eco.NfsB-mCherry)* macrophages (red). In each example, panels include isometric orthogonal yz and xz views corresponding to the xy maximal intensity projection, and indicate the time in minutes from start of movie. Colored arrowheads indicate the conidium Scales as shown. Stills in C,D correspond to Supplementary Movie S5a,c respectively.

### β-glucan is a fungal determinant sufficient for shuttling

Because shuttling was independent of conidial metabolic activity, we hypothesized that the spore-derived signal for shuttling was either from the shape or size of the particle, or was a chemical component of the fungal cell wall such as chitin or β-glucan.

To test whether particle size was sufficient to trigger shuttling, we microinjected 1.7-2.2 μm fluorescent particles (approximating the size of *T. marneffei* and *A. fumigatus* conidia) into the tail somite of 2-3 dpf zebrafish embryos. Although the beads were actively phagocytosed by both neutrophils and macrophages, no shuttling events were observed (~60 hours of imaging, 19 experiments) (Fig 6B). During these experiments, we frequently observed efferocytosis of entire bead-laden neutrophils by macrophages (Supplementary Figure S3, Supplementary Movie S5b). While the neutrophil EGFP fluorescent signal was rapidly lost following engulfment, the bead-conjugated fluorophore signal persisted. These experiments determined that being a particle of a particular size was not sufficient to induce shuttling, and that leukocyte phagocytic behaviour towards inert beads was demonstrably different to their response to fungal conidia. This indicates that shuttling is a conidia-specific behaviour, driven by a chemical signal residing within the condium itself.

The cell wall of fungal conidia is primarily composed of polysaccharides (chitin and glucans) and proteins [28]. The β-glucan class of polysaccharides are a major component of the conidial wall and are highly immunogenenic, so represented a promising candidate shuttling mediator. To test whether β-glucan was sufficient to induce shuttling, we first looked for shuttling of zymosan particles. Zymosan particles are approximately 3 μm in diameter and are a derivative of the *Saccharomyces cerevisiae* cell wall, a rich source of β-glucan glucose polymers. We observed 3 unequivocal shuttles from ~30 hours of imaging over 6 experiments (Fig 6A,C). Because zymosan is predominantly β-glucan, these data suggested that β-glucan may be a spore-derived signal sufficient for shuttling.

To more rigorously test the ability of β-glucan itself to trigger shuttling, we tested whether coating plastic beads in β-glucan conferred on them the ability to be shuttled. While uncoated beads were not shuttled (0 shuttles in 19 experiments), beads coated with β-glucan were shuttled at relatively high frequency (10 shuttles during 22 experiments) (Fig 6B,D). Furthermore, treating β-glucan-coated beads with a mixture of β-glucanase enzymes (including endo/exo-1,3-β-D-glucanase and β-glucosidase) significantly reduced the rate of shuttle ascertainment, from 10 shuttles in 22 experiments to 3 shuttles over 24 experiments (Fig 6B).

As a genetic model, we also tested *Δgel1Δgel7Δcwh41 A. fumigatus* spores, which are deficient in their cell wall β-glucan content due to mutation of their β-1,3-glucanosyltransferase (*gel*) genes (62.6% and 42% of wild type at 37°C, and 50°C, respectively) [29]. The shuttle ascertainment rate for conidia from the mutant strain trended lower compared to wild type *A. fumigatus* (27.3% in wildtype vs. 14.8% in mutant) but this difference was not statistically significant (Fig 6A), likely because the ~50% remaining β-glucan on the conidial cell wall remained sufficient to trigger shuttling.

Collectively, these data support the hypothesis that β-glucan is a fungal wall derived molecule that is sufficient to trigger shuttling signals.

### Shuttling also occurs between mammalian neutrophils and macrophages

Our studies in zebrafish revealed that shuttling was a conserved host response to different species of fungi. We hypothesized that this behaviour might also be conserved between phagocytes from different host species, including higher vertebrates such as mammals.

We tested this hypothesis using an *in vitro* assay. Primary mouse bone marrow neutrophils were preloaded with Alexa Fluor^®^ 488-labelled zymosan added to mouse bone marrow derived macrophages, and imaged over time. The transfer of zymosan particles from living neutrophils to macrophages was observed, in a similar fashion to that observed in the zebrafish *in vivo* model. (Fig 7A,B).

**Figure 7.**
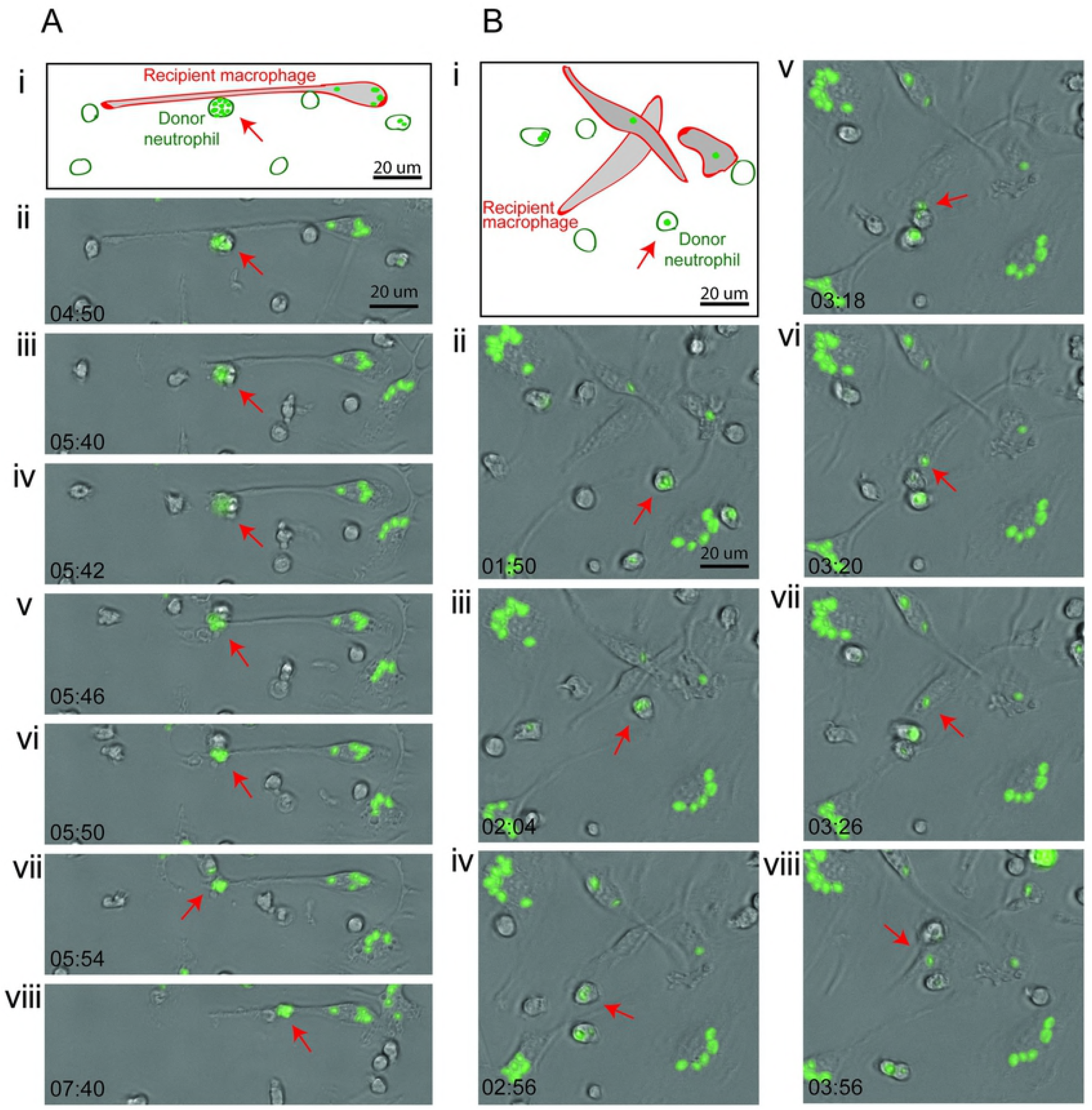
Zymosan shuttles by murine neutrophils and macrophages. A, B. Two sequences demonstrating neutrophil to macrophage shuttling of Alexa Fluor 488-labelled zymosan particle between murine phagocytes *in vitro*. Panel (i) is a schematic showing the elongated, adherent recipient macrophage. Panels (ii-viii) are brightfield photomicrographs with green fluorescence channel overlaid, time points indicated in min:sec. Red arrow indicates the shuttled particle in donor neutrophil (panels ii-vi) and then following shuttling within the recipient macrophage (panels vii-viii). Stills from Supplementary Movie 6.

These data indicate that shuttling is a conserved behaviour of phagocytes in vertebrates from zebrafish to higher mammalian models, and is relevant to host-pathogen interactions during establishment of fungal infections in mammals.

## Discussion

We previously reported the exchange of cytoplasmic fragments from living neutrophils to macrophages during a wound-stimulated inflammatory response [25], although the physiological purpose of this process was unknown. The data presented here reveals that one purpose of cytoplasmic exchange between neutrophils and macrophages is the transfer of phagocytosed microorganisms. It is readily assumed that when a conidium is found within a particular phagocyte during early infection establishment, then it was that particular phagocyte that first phagocytosed it. That is not the case. Conidial shuttling from living neutrophils to macrophages early in fungal infection is an additional and significant aspect of the cell biology of the initial host-pathogen interaction *in vivo*.

Shuttles could only be identified by careful retrospective analysis of live *in vivo* imaging files, which presented a substantial challenge to recognizing them and studying them and their mechanism. From the 188 independent imaging experiments in this report, in total we identified 48 stringently-defined conidial shuttles. Using shuttling ascertainment rates as a surrogate for shuttling incidence was sufficient for comparing conditions when exploring shuttling mechanisms. For example, we observed an overall ascertainment rate of 21.4% (30/140 datasets) for biologic particles (live or dead fungal spores and zymosan particles), compared to 45.5% (10/22) for β-glucan coated beads. From a more biological perspective, for *in vivo* infections following the delivery of 50-100 conidia/inoculum (of which only a minority are initial phagocytosed by neutrophils), the averaged shuttling incidence was at least 19.9% of neutrophil-located spores in the first 3 hr of infection. Shuttling was also sufficiently common for multiple occurrences to be recognized in some movies. Collectively, these observations indicate that shuttling is a consistent, recurring phenomenon during infection establishment, and hence has potential to impact the outcome of the host-pathogen interaction.

To differentiate shuttling from other mechanisms of pathogen entry into macrophages (such as direct phagocytosis, efferocytosis [9], metaforosis/lateral transfer [15], and trogocytosis [16]) it was critical to observe both the cellular origin of shuttled conidium and the moment of transfer. The only possible way to do this was to perform high-resolution 4D confocal microscopy with both high spatial and temporal resolution. Our *in vivo* zebrafish model provided fluorescent labelling clearly distinguishing the two phagocyte lineages, and imaging conditions were optimised for low phototoxicity. While this enabled high spatiotemporal resolution imaging for multiple hours, the imaging volumes often contained considerable biological complexity (high cell densities, cells entering/leaving imaging volume etc.), which made identifying potential interactions quite challenging. It should also be noted that although the total number of inoculated conidia per experiment was only 50-100 particles, only a fraction were within the imaged volume (Fig 4B,C). For these reasons and those mentioned earlier, the shuttling incidence that we report certainly underestimates the absolute rate.

The collective attributes of shuttles distinguish shuttling from all other forms of previously-described conidial transfer between leukocytes. Shuttles were unidirectional (neutrophil to macrophage), occurred only in the first hours after inoculation, and very distinctively, donor neutrophils were alive and mobile before and after shuttling and could shuttle one or more conidia. Recipient macrophages were also alive and mobile, and could be spore-naïve or pre-laden prior to shuttling. Shuttles were preceded by highly regionalized neutrophil-macrophage interactions and occurred through focal cell-to-cell interactions analogous to an intercellular synapse that sometimes resulted in tethering of the two cells together. Macrophages sometimes received aliquots of neutrophil cytoplasm along with the donated spore. This was demonstrated in some cases to be the concomitant transfer of neutrophil membrane around shuttled conidia, consistent with shuttles being the transfer of conidia-laden phagosomes between donor neutrophil and recipient macrophage, rather than just of conidia themselves. This transfer of donor cell membrane also distinguishes shuttling from “non-lytic exocytosis”, as described for the expulsion of previously-phagocytosed *Cryptococcus neoformans* from macrophages [13]. Furthermore, although non-lytic exocytosis expels the pathogen from a macrophage, it has not yet been described in the context of a concurrent interaction with another leukocyte lineage. The cytoplasmic exchange did not, however, provide a durable marker of shuttle occurrence, because the EGFP signal rapidly disappeared, mostly likely due to acidification of the phagolysosome, as is dramatically demonstrated by our example of the efferocytosis of an entire bead-laden neutrophil (Supplementary Fig S3, Supplementary Movie S5).

Both dead and live conidia, labelled with either calcofluor or Alexa Fluor dye, were shuttled. This was consistent with a conidium-directed chemical stimulus driving shuttling, and excluded the possibility that shuttling was a conidium-labelling artefact. Furthermore, shuttling of conidia was conserved between two opportunistic fungal pathogen species, but the kinetics of shuttling was pathogen-specific. This suggested that shuttling was driven by a component common to the conidial cell wall of both species, but one present at different levels or exposed to phagocytes to different degrees [30]. Shuttling of zymosan particles provided further evidence locating a shuttling trigger to the cell wall (Fig 5A,C), leading β-glucan to be identified as a fungal-derived signal sufficient to drive shuttling of plastic beads (Fig 5B,D). A mutant *A. fumigatus* strain with reduced β-glucan trended to lower shuttling rates, also consistent with the hypothesis that conidial β-glucan directly drives shuttling. Since zymosan particles were observed to be shuttled from murine neutrophils to macrophages *in vitro* (Fig 7), shuttling is a conserved phenomenon between vertebrates and, as for other highly conserved phenomena in host defense, likely to play an important role in the outcome of infection.

Our current model for shuttling begins with priming of the spore-laden donor neutrophil and its engagement with the recipient macrophage through pre-shuttle contacts. Within the neutrophil, cytoskeletal rearrangement relocates the conidium within a membrane-lined phagosome towards the side of neutrophil proximate to the recipient macrophage. The conidium, still within its phagosome, is then transferred from the donor neutrophil to the recipient macrophage (Fig 8). The β-glucan-dependent molecular signals involved remain unknown.

**Figure 8.**
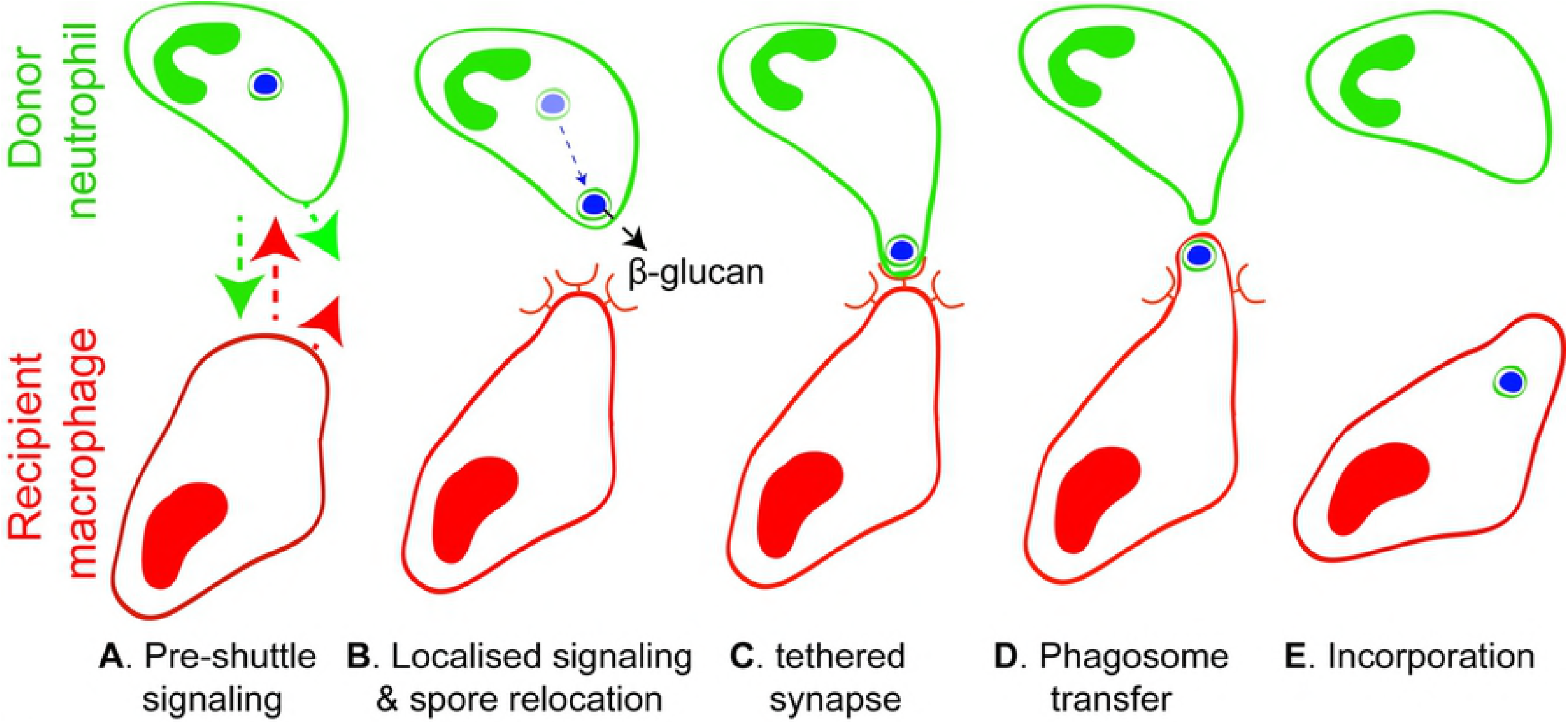
Model of neutrophil to macrophage conidial shuttles. Schematic indicates 5 steps in neutrophil to macrophage conidial shuttling that accommodates morphological and mechanistic insights from these studies. Undefined signals slow the donor neutrophil and recipient macrophage and bring them into proximity (a), leading to β-glucan dependent intercellular shuttling signals and spore relocation within donor cell toward recipient macrophage (b). An intercellular synapse forms with tethering (c), leading to phagosome transfer (d), and it incorporation into recipient macrophages, at least initially retaining components of the membrane-lined donor cell phagosome (e). Both donor and recipient cells remain active following shuttling and eventually both depart (e).

Phagocytosis of fungal pathogens by mammalian leukocytes involves a cluster of pathogen recognition receptors (PRRs) including Dectin-1, Toll-like receptor 2 (TLR2) and Macrophage Mannose Receptor (MMR) [31]. As the mammalian receptor for β-glucan is Dectin-1 [27], it is likely that this receptor and downstream signaling pathways will be involved in conidial shuttling as well as phagocytosis. Although a homolog of mammalian Dectin-1 has yet to be identified in the zebrafish genome, known downstream signaling components such as spleen tyrosine kinase (Syk) have been studied [32]. As the neutrophil is clearly viable following the exchange, and as tethering involves only a small portion of the neutrophil membrane, it is improbable that the triggers are broadly displayed “eat me” signals of imminently apoptotic neutrophils such as phosphatidyl serine or calreticulin [33]. However, regionalized display of such signals might be possible. Testing these hypotheses will be challenging and will require cell-specific and temporally constrained approaches, as their global inhibition will inhibit initial neutrophil phagocytosis of conidia, which is a prerequisite for shuttling.

Neutrophil-to-macrophage pathogen shuttling poses other intriguing mechanistic questions. *Is it unique to fungal infection or does it occur more widely?* Neutrophil-to-macrophage cytoplasm transfer was observed during inflammation [25] suggesting that shuttling may be regulated by inflammatory cytokines. Macrophage cytoplasm transfer to melanoma tumor cells has recently been shown to augment metastatic dissemination, and may be another manifestation of this behaviour [3]. *Is shuttling achieved by repurposing of existing cellular machinery?* The tethering of separating participating cells immediately after the interaction points to potential involvement of the neutrophil uropod, a structure under much traction stress and rich in actively rearranging cytoskeletal components such as actin-myosin bundles [34]. Shuttling may be another manifestation of co-opted trogocytosis mechanisms, as described for macrophage-to-macrophage exchange of Gram negative bacteria [16]. However, trogocytosis-associated intercellular bacterial exchanges cannot involve β-glucan signalling.

The most tantalizing question is: *what is the impact on the microbiological outcome of the infection?* Tied up with this is whether shuttling serves to benefit the host or the pathogen. We recently showed in zebrafish models that macrophages provide an intracellular niche protecting *T. marneffei* conidia from neutrophil fungicidal activity [20]. *A. fumigatus* conidia are also protected by macrophages from neutrophil fungicidal activities [20,23]. Hence fungal-driven shuttling may have evolved to optimize the location of invading conidia into the less hostile intracellular environment of macrophages. Certainly, for these two pathogens, shuttling augments initial conidial redistribution away from the unfavourable neutrophil intracellular environment into their viability-enhancing macrophage intracellular niche. Alternatively, shuttling may be a host-defense mechanism aiding adaptive immunity. Neutrophils are ineffective antigen-presenting cells, whereas macrophages specialize in this, therefore the potential outcome of neutrophil-to-macrophage transfer would be to make pathogen antigens accessible to the adaptive immune system. Delineating the viability outcome for shuttled conidia will require tools for tracing individual shuttled spore fate and/or longitudinal viability throughout the animal (not just in the limited high-magnification imaged volume required to observe its occurrence), and for selectively impairing shuttling but not phagocytosis, neither of which is currently possible *in vivo*, where shuttling is most definitively observed.

Now that this additional phagocyte behaviour during fungal infection establishment has been recognized, its implications must be factored into future understanding of the initial host-pathogen interaction specifically, and into the view of neutrophil and macrophage behaviors generally.

## Materials and Methods

### Zebrafish

Zebrafish strains were wildtype (AB*) carrying single transgenes or combinations of: *Tg(mpx:EGFP)^i113^* [35]; *Tg(mpeg1:Gal4FF)^gl25^* [25]; *Tg(UAS-E1b:Eco.NfsB-mCherry)^c264^* (Zebrafish International Stock Centre, Eugene, OR); *Tg(mpeg1:mCherry-CaaX)^gl26^* [20]; *Tg(mpx:EGFP-CaaX)^gl27^* [20]. Fish were held in the FishCore (Monash University) aquaria using standard practices. Embryos were held at 28°C in egg water (0.06 g/L salt (Red Sea, Sydney, Australia)) or E3 medium (5 mM NaCl, 0.17 mM KCl, 0.33 mM CaCl_2_, 0.33 mM MgSO_4_, equilibrated to pH 7.0); from 12 hpf, 0.003% 1-phenyl-2-thiourea (Sigma-Aldrich) was added. All zebrafish embryos and larvae used in experiments were younger than 7 dpf. Zebrafish exhibit juvenile hermaphroditism, so gender balance in embryonic and larval experiments was not a consideration [36].

### Ethics and Biosafety Statement

All animal experiments followed appropriate NHMRC guidelines. Zebrafish experiments were conducted under protocols approved by Ethics Committees of the Monash University (MAS/2010/18, MARP/2015/094). Zebrafish experiments were performed under Institution Biosafety Committee Notifiable Low Risk Dealing (NLRD) approval PC2-N23-10 (Monash University). *T. marneffei* and *A. fumigatus* were assigned to Risk Group 2 at the time these approvals were granted. In most jurisdictions, including endemic regions, *T. marneffei* is a risk group 2 organism. Protocols for mouse experiments were approved by the Walter and Eliza Hall Institute Animal Ethics Committee.

### *Talaromyces marneffei* and *Aspergillus fumigatus*

The *T. marneffei* strain SPM4 used in this study is a derivative of the FRR2161 type strain [37]. For *A. fumigatus*, wild type *CEA10* [38] and mutant *Δgel1Δgel7Δcwh41* [29] triple mutant strains were used.

To prepare fresh conidia for injection, *T. marneffei* and *A. fumigatus* conidial suspensions were inoculated onto Sabouraud Dextrose (SD) medium and cultured at 25°C for 10-12 days when the cultures were conidiating. Conidia were washed from the plate with dH_2_O, filtered, sedimented (6000 rpm, 10 min), resuspended in dH_2_O and stored at 4°C. For inoculation, conidia were resedimented and resuspended in Phosphate Buffered Saline (PBS). Fungal colony forming unit (CFUs) numbers per embryo were determined as previously described [20].

Cold-inactivation of *T. marneffei* conidia and calcofluor staining was as described previously [20,25]. To inactivate *A. fumigatus* conidia, they were γ-irradiated with 10 kGy [39] from a Gammacell 40 Exactor (Theratronics) as previously described [20] and verified as dead by lack of growth after 5 days incubation. Irradiated conidia still stained well with calcofluor and were microinjected at the same dilution of stock as used for live conidia.

### Zebrafish infection with *Talaromyces marneffei* and *Aspergillus fumigatus*

Freshly-prepared *T. marneffei* and *A. fumigatus* conidia stocks for these experiments were stored at 4°C for <2 months. For inoculation, 52 hpf tricaine-anesthetized embryos were mounted on an agar mould with head/yolk within the well and tail laid flat on the agar. The fungal conidial suspension was inoculated intramuscularly into a somite aligned to the yolk extension tip for local infection [25,40] using a standard microinjection apparatus (Pico-Injector Microinjection System from Harvard Apparatus) and thin wall filament borosilicate glass capillary microinjection needle (SDR Clinical Technology, prepared using a P-2000 micropipette puller, Sutter Instruments). Inoculated embryos were held at 28°C. The delivered conidial dosage was verified by immediate CFU enumeration on a group of injected embryos [20]. It took approximately 10 minutes to commence imaging after inoculation; in this report, the zero time point (t=0) is taken as the beginning of imaging.

### Calcofluor and Alexa Fluor 405 staining of conidia

For calcofluor staining, spores were incubated in 10 mM calcofluor White (Sigma) for 30 minutes, followed by two washing steps and resuspension in distilled water.

To stain fungal conidia with Alexa Fluor 405 NHS Succinimidyl Ester (Life Technologies), 10 μL of Alexa Fluor dye was added to 200 μL of suspended conidia with gentle shaking at room temperature for 30 minutes followed by washing steps with PBS pH 8, 25 mM Tris pH 8.5 and finally resuspended in PBS pH 7, according to the supplier’s protocol.

### Zymosan particles

Zymosan A particles from *Saccharomyces cerevisiae* (Sigma) with average size of 3 μm were stained by calcofluor as for fungal conidia prior to microinjection.

### Plastic Beads

SPHERO^−^ fluorescent Light-Yellow Particles, high Intensity sized 1.7-2.2 μm (SPHEROTECH) (concentration 1.0% w/v in deionized water with 0.02% Sodium Azide) were used. These particles were kept in room temperature. Excitation and emission wavelengths were 400 and 450 nm. Customised commercially-prepared Light-Yellow Particles coated with laminarin as a source of β-glucan (SPHERO^™^ Laminarin Polysaccharide Fluorescent Particles, Lt. Yellow, 1.5-1.99 μm, Catalog no. LPFP1545-2, Lot no. AH01) were also used. Laminarin for coating was from *Laminaria digitata* (primarily poly(β-Glc-[1→3]) with some β- [1→6] interstrand linkages and branch points; Sigma) [41,42].

### *In vitro* studies using murine phagocytes

Primary C57BL/6J mouse bone marrow leukocytes were collected and purified as previously described [43,44]. Macrophages were plated at 5×10^3^ of an 8-well plate incubated in Dulbecco’s modified Eagle’s medium with 10% foetal bovine serum and 20% L-929 conditioned medium for 16 hr. Primary bone marrow neutrophils were pre-loaded with Alexa Fluor 488-labelled opsonized zymosan particles for 1 hr at 37°C in Dulbecco’s modified Eagle’s medium and 10% fetal bovine serum. Preloaded neutrophils were added to adherent macrophages at 10^5^ cells per well. Imaging was performed on a Nikon Biostation IM-Q at 37°C/10% CO_2_.

### Microscopy and image processing

Routine brightfield and fluorescence imaging of zebrafish used an Olympus MVX10 stereo dissecting microscope with MV PLAPO 1X & 2XC objectives fitted with Olympus DP72 camera and Cellsense standard software version 1.11.

Confocal microscopy used a Zeiss LSM 5 Live with a Plan-Apochromat 20x, 0.8 NA objective. ZEN software (2012, black edition 64 bit) was used for acquisition and images were 16-bit 512 x 512 pixels. Z-depth ranged from 35-130 μm (72±23 μm) and composed of 20-40 slices (31±4). Time intervals between z-stacks were set as zero to perform continuous acquisition (z-stack acquisition took 33.24±9.50 seconds). Excitatory laser wavelengths were 405 nm for calcofluor, 489 nm for EGFP and 561 nm for mCherry. Emission detection used a BP495-555 filter for calcofluor and EGFP and a LP575 filter for mCherry. Excitation/emission conditions for Light Yellow particles were the same as for calcofluor.

### Image processing and analysis

All fluorescent image analyses were performed primarily in Imaris (BitPlane) software version 8.1.2 on Venom (Intel^®^ Core™ i7-4770 Processor, 3.4 GHz) or Titan (Intel^®^ Xeon^®^ Processor E5-2680 v2 (2 x 2.80 GHz), RAM: 128 GB) computers (Monash Micro Imaging facility, Monash University). Some analyses used Fiji (ImageJ 1.46r) and Matlab^®^ (The Mathworks™, Natick Mass, USA). For Fig 4a, data were analyzed in the R program using ggplot2 as previously [26,45]. Figures were constructed using Adobe Illustrator CS5 (version 15.0.0).

### Shuttle detection and definition

All *in vivo* shuttles were detected by systematic manual frame-by-frame inspection of movies. For these studies, a “shuttle” was stringently defined as a spore transfer event meeting all of the following criteria: (1) both donor and recipient cells were imaged *in toto* before, during and after the shuttle; (2) both donor and recipient cells demonstrated their viability before and after shuttling by migration; (3) the moment of donor-to-recipient cell transfer was visualised; (4) z-stack viewing unequivocally confirmed that conidia were within donor and recipient cells prior to and after the shuttle. Experience taught that shuttles were most easily recognized by watching movies in reverse and tracing the source of individual macrophage-located conidia. The identity of all 46 unequivocal shuttles meeting these criteria contributing to this report is assigned in Supplementary Figure S1 and Supplementary Table S1, and is indicated throughout the report.

### ShuttleFinder software

The confocal time series were imported into Matlab^®^ (version 8.1.0.604, R2013a) utilising the bioformats toolbox [46] and the conidia were tracked with PhagoSight [27]. The data input consisted of confocal time series with three channels, each containing the fluorescence of one cell type/conidium. PhagoSight was designed to track phagocytes in confocal time series. Since conidia are smaller than phagocytes, the reduction step of PhagoSight was only applied to large files, those that would have taken more than three days to process without it. To reduce the likelihood of false negatives, the automatic determined threshold for background separation by PhagoSight was lowered by 10%. For each file, only the longest tracks were analyzed (upper third of track length over time). PhagoSight was used in command line mode without user interaction to allow for automated processing, using the MASSIVE cluster [47].

PhagoSight calculates a bounding box, which described the volume surrounding each tracked spore for each time frame. The intensity of the voxels in the two channels describing neutrophils and macrophages was summed over this bounding box and a proportional index r between both was calculated

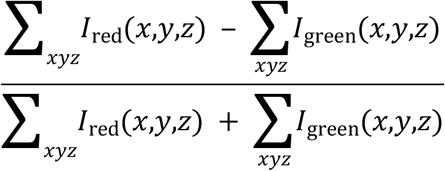

with *I* describing the Intensity of one channel. This ratio was smoothed with a moving average filter over three imaging frames to remove noise caused by imperfections in the tracking process. Subsequently, a point in a track was defined as being in a macrophage (red channel) if the values lie between 1 and 0.2, conversely in a neutrophil (green channel) for a value between −0.2 and −1. To be classified as a candidate shuttle event, the r values for a conidium track had to pass from either −0.2 to 0.2 (for a neutrophil to macrophage shuttle) or vice versa.

The time the tracked conidium reached the threshold was considered the beginning of the shuttle. The end of the shuttle event was defined by the track leaving the threshold area.

### Statistics

Descriptive and analytical statistics were prepared in Prism 5.0c (GraphPad Software Inc). Unless otherwise stated, data are mean±SD, with p-values generated from two-tailed unpaired t-tests for normally distributed continuous variables, and Chi-squared tests for categorical variables.

## Acknowledgements

We thank our colleagues who have provided comment and advice on this project and/or encouraged us to persist as it progressed over several years, particularly: Dr M.C. Keightley, Dr P. Currie. This work was supported by the Multi-modal Australian ScienceS Imaging and Visualisation Environment (MASSIVE) (www.massive.org.au).

## Grant support

GL, AA, National Health and Medical Research Council (461208, 637394, 1044754, 1069284).

BC, National Health and Medical Research Council (637367) Australian Research Council (DP1094854), National Institutes of Health (5RO1HL124209).

FE and LP, Australian Postgraduate Award, Walter and Eliza Hall Institute Edith Moffatt Scholarship.

FE, Monash University Bridging Postdoctoral Fellowship, and while working on writing this manuscript while in the laboratory of Dr D. Irimia, he was supported by the MGH BioMEMS Center (National Institutes of Health, Grant EB002503).

VP, Monash Graduate Scholarship, Monash International Postgraduate Research Fellowship, Monash Postgraduate Publication Award.

JO, Dora Lush Scholarship (National Health and Medical Research Council).

The Australian Regenerative Medicine Institute is supported by grants from the State Government of Victoria and the Australian Government.

## Author Contributions

Listed using the CRediT taxonomy.

## Conceptualization

Vahid Pazhakh, Felix Ellett, Luke Pase, Joanne A. O’Donnell, Ben A. Croker, Alex Andrianopoulos, Graham J. Lieschke.

## Formal analysis

Vahid Pazhakh, Felix Ellett, Joanne A. O’Donnell, Keith E. Schulze, R. Stefan Greulich, Ben A. Croker, Alex Andrianopoulos, Graham J. Lieschke.

## Funding acquisition

Alex Andrianopoulos, Graham J. Lieschke.

## Investigation

Vahid Pazhakh, Felix Ellett, Joanne A. O’Donnell, R. Stefan Greulich, Ben A. Croker, Graham J. Lieschke.

## Methodology

Vahid Pazhakh, Felix Ellett, Joanne A. O’Donnell, R. Stefan Greulich, Keith Schulze, C. Carlos Reyes-Aldasoro, Ben A. Croker, Alex Andrianopoulos, Graham J. Lieschke.

## Software

R. Stefan Greulich, C. Carlos Reyes-Aldasoro.

## Supervision

Ben A. Croker, Alex Andrianopoulos, Graham J. Lieschke.

## Writing – original draft

Vahid Pazhakh, Felix Ellett, Alex Andrianopoulos, Graham J. Lieschke.

## Writing – review and editing

Vahid Pazhakh, Felix Ellett, Luke Pase, Keith Schulze, Joanne A. O’Donnell, R. Stefan Greulich, C. Carlos Reyes-Aldasoro, Ben A. Croker, Alex Andrianopoulos, Graham J. Lieschke.

## SUPPLEMENTARY FIGURE LEGENDS

**Supplementary Figure S1. Imaging datasets.** Details of the imaging datasets in which the defining set of shuttles of 13 *T. marneffei* (A) and 7 *A. fumigatus* (B) conidia meeting stringent definition criteria were found. Graphs show the distribution of imaging file lengths, which files contained a shuttle (black columns), the shuttle ID (#), the shuttle movie length (L) and the time of shuttle (yellow mark in black column, and red numeral in minutes). Corresponds to Figs 1, 2, 4 and Supplementary Table S1.

**Supplementary Figure S2. Examples of probable shuttles** A variety of shuttles of conidia or particles (blue) from *Tg(mpx:EGFP)* neutrophils (green) to *Tg(mpeg1:Gal4FF)x(UAS-E1b:Eco.NfsB-mCherry)* macrophages (red). In each example, panels include isometric orthogonal yz and xz views corresponding to the xy maximal intensity projection, and indicate the time in minutes from start of movie. Colored arrowheads indicate the conidium/particle within donor neutrophil (green), at the point of intercellular transfer (white) and in the recipient macrophage (red). **A-C. Probable shuttles of conidia.** A. A probable shuttle in which the condium is not clearly resolved as fully contained within the donor neutrophil. B-C. Probable shuttles of conidia in which the point of cell-to-cell contact is not clearly displayed. D. An example of a crowded field with multiple neutrophils and macrophages in which initially there are neutrophils laden with conidia and by the end conidia are mostly within macrophages, although the transfer of conidia is not clearly seen. Scales as shown. Stills in A-D correspond to Supplementary Movie S4a-d respectively.

**Supplementary Figure S3. Efferocytosis of an entire bead-laden neutrophil.** Phagocytosis of inert 2 μm plastic beads (blue) by *Tg(mpx:EGFP)* neutrophils (green), followed by efferocytosis of the whole particle-laden neutrophil by a *Tg(mpeg1:Gal4FF)x(UAS-E1b:Eco.NfsB-mCherry)* macrophage (red). Subsequently the EGFP signal of the engulfed neutrophil is extinguished although the Alexa Fluor signal (blue) of the plastic beads persists (right panel). Panels include isometric orthogonal yz and xz views corresponding to the xy maximal intensity projection, and indicate the time in minutes from start of movie. White arrowheads follow the neutrophil of interest through the process. Scale as shown. Stills from Supplementary Movie S5b.

## SUPPLEMENTARY MOVIE LEGENDS

**Supplementary Movie S1. Six examples of live *T. marneffei* conidial shuttles.** Shuttles are of live calcofluor-stained conidia (blue) from a *Tg(mpx:EGFP)* neutrophil (green) to a *Tg(mpeg1:Gal4FF)x(UAS-E1b:Eco.NfsB-mCherry)* macrophage (red). Movies run in series. Movies are paused at the moment of shuttling, with the point of transfer labelled (white arrow). a. Standard shuttle (corresponds to Fig 1A). b. Standard shuttle with tethered recipient macrophage (corresponds to Fig 1E). c. Standard shuttle with tethered donor neutrophil (corresponds to Fig 2A). d. Standard shuttle with tethered departing donor neutrophil. e. Standard shuttle of multiple spores in quick succession (corresponds to Fig 2B). f. Two conidia shuttled asynchronously (corresponds to Fig 2C).

**Supplementary Movie S2. Six examples of *A. fumigatus* conidial shuttles.** Shuttles are of live calcofluor-stained conidia (blue) from a *Tg(mpx:EGFP)* neutrophil (green) to a *Tg(mpeg1:Gal4FF)x(UAS-E1b:Eco.NfsB-mCherry)* macrophage (red). Movies run in series. Movies are paused at the moment of shuttling, with the point of transfer labelled (white arrow). a. Standard shuttle. b. Standard shuttle. c. Standard shuttle of Alexa Fluor 405-stained conidium (corresponds to Fig 2E). d. Shuttling involving a highly polarized and tethered neutrophil and macrophage interaction. e. Two independent shuttles occurring in the same field. f. Two conidia shuttled together.

**Supplementary Movie S3. Four examples of dead *T. marneffei* conidial shuttles.** Shuttles are of dead calcofluor-stained conidia (blue) from a *Tg(mpx:EGFPCaaX)* neutrophil (green) to a *Tg(mpeg1:mCherryCaaX)* macrophage (red). Movies run in series. Movies are paused at the moment of shuttling, with the point of transfer labelled (white arrow). These reporter lines localize the fluorophore to the membrane, enabling these movies to display volume-rendered version of donor neutrophil and recipient macrophages in parallel (right panels). a-c. Three independent shuttles of individual dead conidia, all occurring in the same movie (shuttle (a) corresponds to Fig 3A). d. Standard shuttle of a dead conidium.

**Supplementary Movie S4. Four examples of probable conidial shuttles not meeting all definition criteria.** Shuttles are of live calcofluor-stained conidia (blue) from a *Tg(mpx:EGFP)* neutrophil (green) to a *Tg(mpeg1:Gal4FF)x(UAS-E1b:Eco.NfsB-mCherry)* macrophage (red). Movies run in series. Movies are paused at the moment of shuttling, with the point of transfer labelled (white arrow). a. Probably shuttle in which the conidium is not unequivocally resolved as being within the donor neutrophil (corresponds with Supplementary Fig S2A). b,c. Probable shuttles in which direct intercellular contact between donor neutrophil and recipient macrophage is not clearly displayed (corresponds with Supplementary Fig S2B-C). d. Crowded field in which there are initially neutrophil-laden conidia, and at the end, macrophage-laden conidia, but the crowding obscures probable conidial shuttling (corresponds with Supplementary Fig S2D).

**Supplementary Movie S5. Examples of shuttles of non-conidial particles.** Shuttles are of live calcofluor-stained conidia (blue) from a *Tg(mpx:EGFP)* neutrophil (green) to a *Tg(mpeg1:Gal4FF)x(UAS-E1b:Eco.NfsB-mCherry)* macrophage (red). Movies run in series. Movies are paused at the moment of shuttling, with the point of transfer labelled (white arrow). a. Shuttle of zymosan particle (corresponds to Fig 6C). b. Efferocytosis (not a shuttle) of whole neutrophil laden with plastic beads (corresponds to Supplementary Fig S3). c. Shuttle of β-glucan coated plastic beads (corresponds to Fig 6D).

**Supplementary Movie S6. Two examples of zymosan shuttles between murine neutrophils and macrophages *in vitro*.** Shuttles of zymosan particles between murine neutrophils preloaded with Alexa Fluor 488-labelled zymosan and adherent murine macrophages in an *in vitro* assay. Photomicrographs are brightfield views overlaid with green fluorescence channel. White arrows in paused frames indicate the shuttle. Time stamps are provided in the corresponding Fig 7 stills. a. Shuttle (arrowed) (corresponds with Fig 7A). b. Shuttle (arrowed) (corresponds with Fig 7B).

## References

1. Kaufmann SH (2008) Immunology’s foundation: the 100-year anniversary of the Nobel Prize to Paul Ehrlich and Elie Metchnikoff. Nat Immunol 9: 705–712.

2. Uribe-Querol E, Rosales C (2017) Control of Phagocytosis by Microbial Pathogens. Frontiers in Immunology 8.

3. Roh-Johnson M, Shah AN, Stonick JA, Poudel KR, Kargl J, et al. (2017) Macrophage-Dependent Cytoplasmic Transfer during Melanoma Invasion In Vivo. Developmental cell 43: 549–562 e546.

4. McCoy-Simandle K, Hanna SJ, Cox D (2016) Exosomes and nanotubes: Control of immune cell communication. Int J Biochem Cell Biol 71: 44–54.

5. Weidle UH, Birzele F, Kollmorgen G, Ruger R (2017) The Multiple Roles of Exosomes in Metastasis. Cancer Genomics Proteomics 14: 1–15.

6. Zomer A, Maynard C, Verweij FJ, Kamermans A, Schafer R, et al. (2015) In Vivo imaging reveals extracellular vesicle-mediated phenocopying of metastatic behavior. Cell 161: 1046–1057.

7. Spinner JL, Winfree S, Starr T, Shannon JG, Nair V, et al. (2014) Yersinia pestis survival and replication within human neutrophil phagosomes and uptake of infected neutrophils by macrophages. J Leukoc Biol 95: 389–398.

8. Martin CJ, Peters KN, Behar SM (2014) Macrophages clean up: efferocytosis and microbial control. Curr Opin Microbiol 17: 17–23.

9. Karaji N, Sattentau QJ (2017) Efferocytosis of Pathogen-Infected Cells. Frontiers in immunology 8: 1863.

10. Yang CT, Cambier CJ, Davis JM, Hall CJ, Crosier PS, et al. (2012) Neutrophils exert protection in the early tuberculous granuloma by oxidative killing of mycobacteria phagocytosed from infected macrophages. Cell host & microbe 12: 301–312.

11. Bain JM, Lewis LE, Okai B, Quinn J, Gow NA, et al. (2012) Non-lytic expulsion/exocytosis of Candida albicans from macrophages. Fungal Genet Biol 49: 677–678.

12. Nicola AM, Robertson EJ, Albuquerque P, Derengowski Lda S, Casadevall A (2011) Nonlytic exocytosis of Cryptococcus neoformans from macrophages occurs in vivo and is influenced by phagosomal pH. MBio 2.

13. Ma H, Croudace JE, Lammas DA, May RC (2007) Direct cell-to-cell spread of a pathogenic yeast. BMC Immunol 8: 15.

14. Shah A, Kannambath S, Herbst S, Rogers A, Soresi S, et al. (2016) Calcineurin Orchestrates Lateral Transfer of Aspergillus fumigatus during Macrophage Cell Death. Am J Resp Crit Care 194: 1127–1139.

15. Armstrong-James D, de Boer L, Bercusson A, Shah A (2018) From phagocytosis to metaforosis: Calcineurin’s deadly role in innate processing of fungi. PLoS pathogens 14: e1006627.

16. Steele S, Radlinski L, Taft-Benz S, Brunton J, Kawula TH (2016) Trogocytosis-associated cell to cell spread of intracellular bacterial pathogens. eLife 5.

17. Kaufmann SHE, Dorhoi A, Hotchkiss RS, Bartenschlager R (2018) Host-directed therapies for bacterial and viral infections. Nat Rev Drug Discov 17: 35–56.

18. Keightley MC, Wang CH, Pazhakh V, Lieschke GJ (2014) Delineating the roles of neutrophils and macrophages in zebrafish regeneration models. The international journal of biochemistry & cell biology 56: 92–106.

19. Renshaw SA, Trede NS (2012) A model 450 million years in the making: zebrafish and vertebrate immunity. Disease Models & Mechanisms 5: 38–47.

20. Ellett F, Pazhakh V, Pase L, Benard EL, Weerasinghe H, et al. (2018) Macrophages protect Talaromyces marneffei conidia from myeloperoxidase-dependent neutrophil fungicidal activity during infection establishment in vivo. PLoS pathogens 14: e1007063.

21. Knox BP, Huttenlocher A, Keller NP (2017) Real-time visualization of immune cell clearance of Aspergillus fumigatus spores and hyphae. Fungal Genet Biol 105: 52–54.

22. Knox DM (1999) Core body temperature, skin temperature, and interface pressure. Relationship to skin integrity in nursing home residents. Adv Wound Care 12: 246–252.

23. Rosowski EE, Raffa N, Knox BP, Golenberg N, Keller NP, et al. (2018) Macrophages inhibit Aspergillus fumigatus germination and neutrophil-mediated fungal killing. PLoS pathogens 14: e1007229.

24. Knox BP, Deng Q, Rood M, Eickhoff JC, Keller NP, et al. (2014) Distinct innate immune phagocyte responses to Aspergillus fumigatus conidia and hyphae in zebrafish larvae. Eukaryotic cell 13: 1266–1277.

25. Ellett F, Pase L, Hayman JW, Andrianopoulos A, Lieschke GJ (2011) mpeg1 promoter transgenes direct macrophage-lineage expression in zebrafish. Blood 117: e49–56.

26. Lammermann T, Afonso PV, Angermann BR, Wang JM, Kastenmuller W, et al. (2013) Neutrophil swarms require LTB4 and integrins at sites of cell death in vivo. Nature 498: 371–375.

27. Henry KM, Pase L, Ramos-Lopez CF, Lieschke GJ, Renshaw SA, et al. (2013) PhagoSight: an open-source MATLAB(R) package for the analysis of fluorescent neutrophil and macrophage migration in a zebrafish model. PloS one 8: e72636.

28. Free SJ (2013) Fungal cell wall organization and biosynthesis. Advances in genetics 81: 33–82.

29. Zhao W, Li CL, Liang JN, Sun SF (2014) The Aspergillus fumigatus beta-1,3-glucanosyltransferase Gel7 plays a compensatory role in maintaining cell wall integrity under stress conditions. Glycobiology 24: 418–427.

30. Arana DM, Prieto D, Roman E, Nombela C, Alonso-Monge R, et al. (2009) The role of the cell wall in fungal pathogenesis. Microbial biotechnology 2: 308–320.

31. Inoue M, Shinohara ML (2014) Clustering of pattern recognition receptors for fungal detection. PLoS Pathog 10: e1003873.

32. Christie TL, Carter A, Rollins EL, Childs SJ (2010) Syk and Zap-70 function redundantly to promote angioblast migration. Developmental biology 340: 22–29.

33. Duffin R, Leitch AE, Fox S, Haslett C, Rossi AG (2010) Targeting granulocyte apoptosis: mechanisms, models, and therapies. Immunol Rev 236: 28–40.

34. Yoo SK, Lam PY, Eichelberg MR, Zasadil L, Bement WM, et al. (2012) The role of microtubules in neutrophil polarity and migration in live zebrafish. Journal of Cell Science 125: 5702–5710.

35. Renshaw SA, Loynes CA, Trushell DM, Elworthy S, Ingham PW, et al. (2006) A transgenic zebrafish model of neutrophilic inflammation. Blood 108: 3976–3978.

36. Hsiao CD, Tsai HJ (2003) Transgenic zebrafish with fluorescent germ cell: a useful tool to visualize germ cell proliferation and juvenile hermaphroditism in vivo. Dev Biol 262: 313–323.

37. Borneman AR, Hynes MJ, Andrianopoulos A (2001) An STE12 homolog from the asexual, dimorphic fungus Penicillium marneffei complements the defect in sexual development of an Aspergillus nidulans steA mutant. Genetics 157: 1003–1014.

38. Fedorova ND, Khaldi N, Joardar VS, Maiti R, Amedeo P, et al. (2008) Genomic islands in the pathogenic filamentous fungus Aspergillus fumigatus. PLoS genetics 4: e1000046.

39. Gumus T, Geegel U, Demirci AS, Arici M (2008) Effects of gamma irradiation on two heat resistant moulds: Aspergillus fumigatus and Paecilomyces variotii isolated from margarine. Radiat Phys Chem 77: 680–683.

40. Benard EL, van der Sar AM, Ellett F, Lieschke GJ, Spaink HP, et al. (2012) Infection of zebrafish embryos with intracellular bacterial pathogens. J Vis Exp 3781 [pii] 10.3791/3781.

41. Brown GD, Gordon S (2001) Immune recognition. A new receptor for beta-glucans. Nature 413: 36–37.

42. Rubin-Bejerano I, Abeijon C, Magnelli P, Grisafi P, Fink GR (2007) Phagocytosis by human neutrophils is stimulated by a unique fungal cell wall component. Cell host & microbe 2: 55–67.

43. Croker BA, Lewis RS, Babon JJ, Mintern JD, Jenne DE, et al. (2011) Neutrophils require SHP1 to regulate IL-1beta production and prevent inflammatory skin disease. Journal of immunology 186: 1131–1139.

44. Masters SL, Gerlic M, Metcalf D, Preston S, Pellegrini M, et al. (2012) NLRP1 inflammasome activation induces pyroptosis of hematopoietic progenitor cells. Immunity 37: 1009–1023.

45. Bhuiyan MS, Ellett F, Murray GL, Kostoulias X, Cerqueira GM, et al. (2016) Acinetobacter baumannii phenylacetic acid metabolism influences infection outcome through a direct effect on neutrophil chemotaxis. Proceedings of the National Academy of Sciences of the United States of America 113: 9599–9604.

46. Linkert M, Rueden CT, Allan C, Burel JM, Moore W, et al. (2010) Metadata matters: access to image data in the real world. Journal of Cell Biology 189: 777–782.

47. Goscinski WJ, McIntosh P, Felzmann U, Maksimenko A, Hall CJ, et al. (2014) The multi-modal Australian ScienceS Imaging and Visualization Environment (MASSIVE) high performance computing infrastructure: applications in neuroscience and neuroinformatics research. Front Neuroinform 8.

